# Kinetics and optimality of influenza A virus locomotion

**DOI:** 10.1101/2024.05.06.592729

**Authors:** Siddhansh Agarwal, Boris Veytsman, Daniel A. Fletcher, Greg Huber

## Abstract

Influenza A viruses (IAVs) must navigate through a dense extracellular mucus to infect airway epithelial cells. The mucous layer, composed of glycosylated biopolymers (mucins), presents sialic acid that binds to ligands on the viral envelope and can be irreversibly cleaved by viral enzymes. It was recently discovered that filamentous IAVs exhibit directed persistent motion along their long axis on sialic acid-coated surfaces. This study demonstrates through stochastic simulations and mean-field theory, how IAVs harness a ‘burnt-bridge’ Brownian ratchet mechanism for directed persistent translational motion. Importantly, our analysis reveals that equilibrium features of the system primarily control the dynamics, even out-of-equilibrium, and that ligand asymmetry allows for more robust directed transport. We show viruses occupy the optimal parameter range (‘Goldilocks zone’) for efficient mucous transport, possibly due to the evolutionary adaptation of enzyme kinetics. Our findings suggest novel therapeutic targets and provide insight into possible mechanisms of zoonotic transmission.

## Introduction

The onset of many infectious diseases hinges crucially on the ability of invading pathogens to traverse the extracellular environment on the way to the host cell membrane [1, 2]. Mucotropic viruses such as influenza A must overcome the extracellular mucosal barrier protecting airway epithelial cells [3]. Despite extensive studies on the biochemistry of virus-host interactions [4, 5] and the lateral diffusion dynamics of viral particles on model membrane systems [6, 7], a quantitative understanding of how viruses move through material barriers remains incomplete.

The mucous layer coating respiratory epithelia with a thickness ranging from ~ 1 µm to 10 µm [8–10] poses a critical barrier for influenza A viruses (IAVs) to overcome. Mucins have abundant binding sites for the IAV surface protein hemagglutinin (HA), effectively immobilizing viruses within the layer. Coordinated beating of cilia on epithelial cells promotes a net mucous flow resulting in clearance of pathogens before they can reach the vulnerable substrate [11–14]. With the estimated mucociliary clearance rates of ~ 1 cm*/*min [15, 16], IAVs have a time window of just *t*_*c*_ ~ 20 min [17] before being cleared from the airway. Clinical isolates of IAVs typically have elongated, filamentous morphologies with a cylindrical diameter between 70 nm to 90 nm and a broad length distribution ranging from 100 nm to over 5 µm [18–20]. Shorter filamentous strands with axial lengths under 300 nm are thought to be the primary vectors spreading infection between cells [17, 21, 22]. For a virion with these dimensions, the timescale to move a distance of ~ 10 µm based on passive diffusion alone is *t*_*d*_ ~ 2 h [17] (see Supplementary Material (SM) for further details). Thus the fraction of IAVs that overcome the mucous barrier is ~ exp(*t*_*d*_*/t*_*c*_) = 2.5 × 10^−3^. This is not the only defense mechanism of the host: epithelial cell surfaces in the periciliary layer beneath the layer of mucus have long, densely packed glycoprotein “brush” elements creating a steric barrier that viruses must penetrate [23, 24], further reducing the number of viruses that can reach the airway epithelium. Therefore viruses with purely diffusive motion have little chance to penetrate the mucociliary layer before being cleared away.

Vahey and Fletcher [17] discovered a key strategy used by IAVs to navigate host barriers: persistent movement powered by its local environment. Optical microscopy of filamentous IAVs revealed that the distribution of the proteins HA and Neuraminidase (NA) on the viral surface is polarized: NA enzyme clusters preferentially associated with the genome-containing end. Experiments on coverslips functionalized with sialic acid (SA) showed persistent translational motion of the virions away from the NA-rich pole, operating via a ‘burnt-bridge’ Brownian ratchet mechanism [25]. Recent work [26] has revealed that this motion is modulated by the HA/NA activity and their spatial organization, with observed differences in transport speeds and gaits. In contrast, influenza C viruses exhibit rolling motion on SA-functionalized surfaces due to the coordinated SA binding/cleavage cycle of their hemagglutininesterase-fusion glycoprotein [7, 27], which imposes a uniform surface distribution of binders & cleavers. For IAVs, the elongated morphology and spatial segregation of HA and NA likely favor translational over rotational modes within native viscoelastic mucinous networks, where pore sizes (~ 100 nm) [28] impose constraints on large-scale rotational movement.

This work examines the translational motion of short filamentous IAVs, with a focus on the “delicate balance” of HA/NA activities hypothesized as a key factor governing their movement [29]. We demonstrate that in our model, due to the irreversible SA-cleaving process, the virions move with constant speed, and this speed exhibits a maximum within a specific range of HA and NA parameter values approximating those of real viruses. This suggests that evolutionary pressures may have selected for viruses optimized to enhance their ability to navigate and penetrate host barriers.

## Model

We develop a simplified 1D representation of the virion’s movement that focuses on its interactions with local SA receptors, capturing key aspects of its trajectory while maintaining computational feasibility. We consider a coarse-grained representation of an IAV as a rigid, 1D rod of length *L* (~ 300 nm). The viral envelope is populated by a fixed, non-diffusive array of HA spikes binding to SA moieties, and NA enzyme complexes cleaving these receptors. Typical mean separations between HA on the IAV envelope are *l*_0_ ~ 10 nm (*l*_0_ ≪ *L*) [30]. Thus, we assume there are 30 ligands distributed uniformly on the simulated virus surface.

We model the virus as positioned at a fixed elevation above a 1D substrate uniformly decorated with SA at an average spacing *l* ~ 2.5 nm reflecting typical glycoprotein densities [17]. HA binding to SA is described as a bimolecular reaction with the off-rate (*k*_off_) from empirical measurements [31], and the on-rate (*k*_on_) estimated by modeling HA trimers and SA as confined complexes following previous studies [17, 26]. We assume each SA receptor can bind to only one HA at a time. NA irreversibly cleaves free SA via first-order kinetics with a catalytic rate *k*_*c*_ [32], in parallel. In our model, HA-SA interactions are represented by Hookean springs with spring constant *S*, which exert forces on the virion. This accounts for glycoprotein linker flexibility [27], coarse-graining fast linker fluctuations while capturing effective entropic elasticity. This assumption is justified by the much faster timescale of NA-SA interactions compared to HA-SA bonds. In the following, we choose *l*_0_ and *k*_on_ as the length and inverse-time scales, respectively, with specific parameter values provided in the SM.

We scale the rates of bond formations and cleaving by exp(−*u/*(*k*_*B*_*T*)) = exp(−Δ*x*^2^*/α*^2^), where *u* is the potential energy of the spring stretched by the value Δ*x*, and 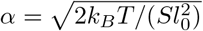 is the non-dimensional thermal fluctuation parameter. The dimensionless binding/unbinding rates *q*_off_ = *K*_*D*_*/*(1 + exp(−Δ*x*^2^*/α*^2^)), *q*_on_ = exp(−Δ*x*^2^*/α*^2^)*/*(1 + exp(−Δ*x*^2^*/α*^2^) are given by transition-state theory (see, e.g., [33]), defining an effective stretch-dependent dissociation constant *q*_off_*/q*_on_ = *K*_*D*_ exp(Δ*x*^2^*/α*^2^), where *K*_*D*_ = *k*_off_*/k*_on_. This approach treats the unstretched state as the reference transition state, modifying rates based on bond stretch without specifying a fixed barrier location [34]. The dimensionless NA cleavage rate is *q*_*c*_ = *K*_*C*_ exp(−Δ*x*^2^*/α*^2^), where *K*_*C*_ = *k*_*c*_*/k*_on_. The exponential dependence on bond stretch represents a simplified model of distance penalty, where SA receptors further from the virus surface are less accessible to NA, thus less likely to be cleaved. In the simulations we use overdamped dynamics for the speed *v* of the virus at position *x*(*t*):

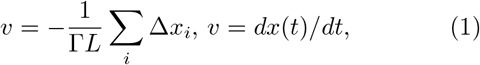

where Γ = *γk*_on_*/S* is a dimensionless hydrodynamic friction parameter representing the ratio of two characteristic timescales: the time for the virus to move under spring forces and the time for bond formation. Here *γ* is the translational friction coefficient scaled by the length *L*. We assume constant virion size orthogonal to the direction of motion. In contrast to conventional Langevin dynamics, Brownian motion is omitted in our analysis, since 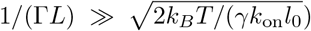, indicating that the spring forces predominate over higher-frequency thermal fluctuations. At each time step, we first stochastically update the configuration of bound springs and cleaved receptors via Gillespie algorithm, an efficient kinetic Monte Carlo approach [35, 36]. Then the net instantaneous force is computed and used to update the velocity and position of the virus in (1).

This insight, that the most important process in the system is the formation, arrangement and cleavage of the bonds, is the key for the analytical treatment. We assume the large separation of timescales between binding/unbinding, on one hand, and cleavage together with virus motion, on another hand. Thus at any time and location, the configuration of bonds is close to the equilibrium one, and we calculate its Gibbs free energy *G* summing the logs of partition functions of bonds formed at each virus site (see the SM for details). When the number of virus sites is large, we can use the continuous limit, and the sum becomes the integral,

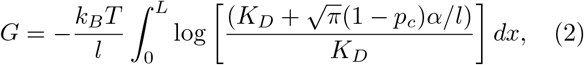

where *p*_*c*_(*x*) is the probability that a substrate site is burned. Here *x* is the coordinate in the co-moving frame of the virus. If the virus moves with a constant speed *v*, the cleaving probability profile *p*_*c*_(*t, x*) in the co-moving frame satisfies the quasi-steady equation,

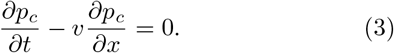

Differentiating equation (2) by time and using equation (3), we can calculate the production of free energy *dG/dt*.

Free energy produced by the irreversible cleaving dissipates through the hydrodynamic friction of the virus as a whole and linker friction due to bonds “snapping” [37]. Equating free-energy production to dissipation when the virus moves with a steady speed *v*, we get

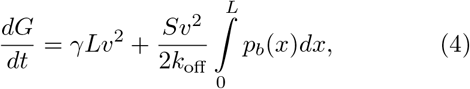

where 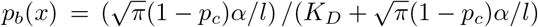 is the binding probability. Calculating the *dG/dt* and the intergral over *p*_*b*_, we get a closed equation for *v*. It can be solved analytically in two limiting cases (see SM): when 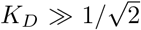 and when 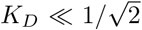. In the first

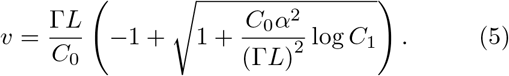

where 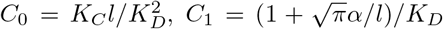. In the second case

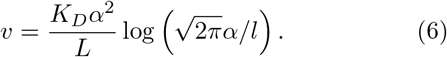

## Results and Discussion

For simplicity, we discuss only two configurations of binders and cleavers on the viral envelope: (i) a uniform distribution of HA and NA over the virion’s surface, and (ii) a spatially segregated case where NA is localized to half of the virion, while HA is uniformly distributed. For the uniform case (Fig. 2A–C), increasing viral length *L* triggers a spontaneous symmetry breaking (SSB) transition from diffusive motion (purple, *L* = 2) to ballistic, directional motion (red, *L* = 30). The mean-squared displacement (MSD) vs time (Fig. 2B) clearly shows this transition: the long-time MSD exponent approaches 2 as *L* increases. NA’s stochastic cleavage activity generates an asymmetric SA receptor concentration profile, inducing spatial anisotropy in HA-SA bond formation probability. This asymmetry results in a net force propelling the virion away from cleaved regions. Consequently, NA-induced SA gradients rectify the stochastic HA-SA bond kinetics into directed motion, creating a self-reinforcing molecular ratchet.

**FIG. 1.**
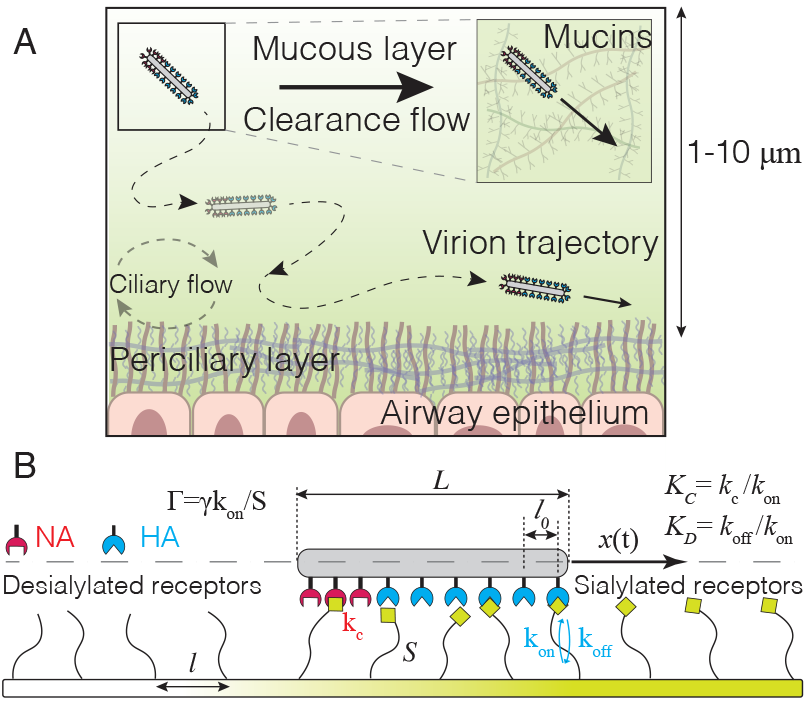
(A) A virion (not to scale) navigates the complex heterogeneous and flowing mucous layer (1–10 µm) composed of highly glycosylated, entangled mucin fibers. (B) Coarse-grained model of a virion as a rigid elongated shell, positioned at a constant offset above a 1D substrate consisting of uniformly coated glycoproteins decorated with SA receptors. HA can form reversible bonds with SA on the surface, while NA cleaves these receptors irreversibly. An HA–SA bond is modeled as a Hookean spring with stiffness *S*, modeling the finite extensibility of the glycoproteins.

**FIG. 2.**
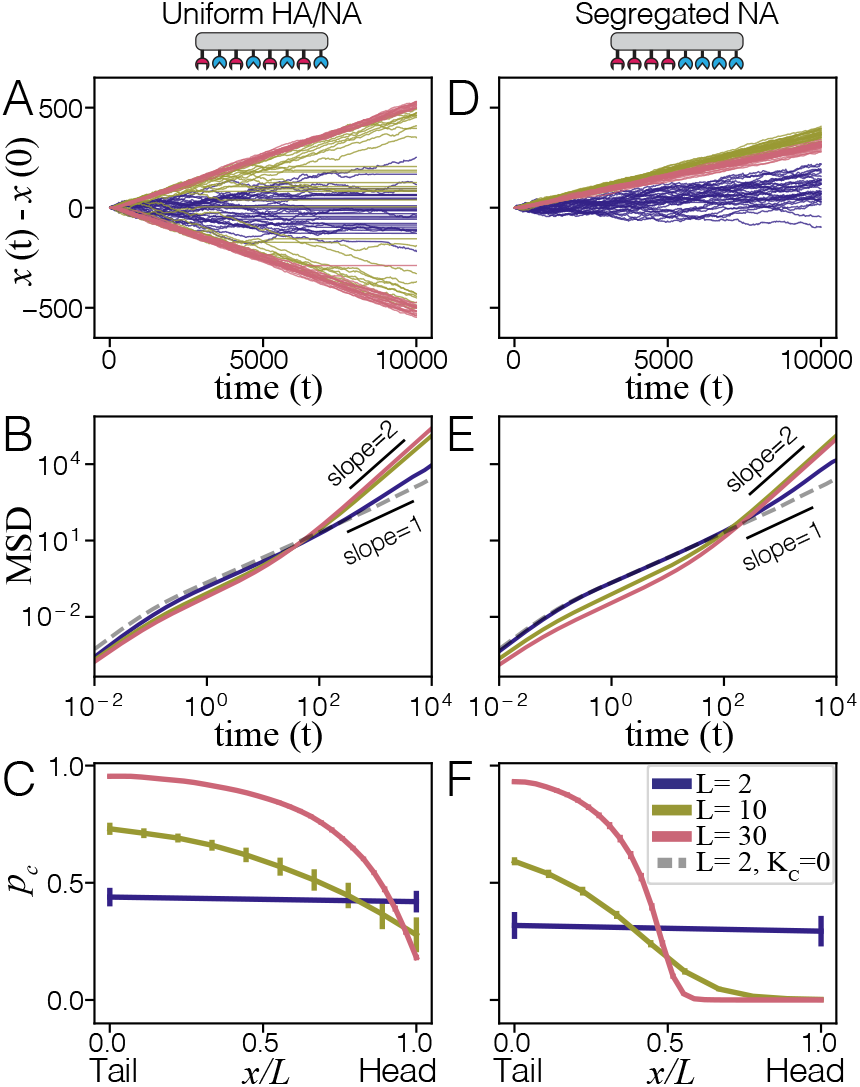
IAV locomotion with uniform (A-C) versus segregated (D-F) HA/NA distributions for different lengths *L*. Displacement from the initial position *x*(*t*) − *x*(0) plotted against time *t* demonstrates SSB for uniform HA/NA distribution (A), while the virus consistently moves away from the NA pole for the segregated case (D). (B, E) Mean-squared displacement (MSD) versus time plots show the transition from diffusive (slope = 1) to ballistic motion (slope = 2) as *L* increases. (C, F) The averaged SA cleavage probabilities along the virus *x/L* highlight the distinct SA cleavage profiles and consequent directional biases between the two distributions.

In the segregated NA case (Fig. 2D–F) the symmetry is absent, and the virion robustly propels away from the NA-rich side at any *L*. The diffusive, non-cleaving case (*L* = 2, *K*_*C*_ = 0, dashed gray) is shown for reference. Analysis of steady-state cleaved SA profiles beneath the virus (Fig. 2C, F) reveals the mechanisms underlying ballistic motion. Both uniform and segregated HA/NA configurations exhibit an *L*_*c*_ ~ 10 for ballistic motion onset. The uniform case spontaneously develops a head-tail gradient, while the segregated case inherently generates an SA concentration gradient due to asymmetric NA distribution. We note that the viruses in our model resemble active Brownian particles that move with constant speed, slowly changing direction or turning when bumping into obstacles [38, 39].

Analytical theory provides a quantitative framework to understand the steady-state cleaving profile *p*_*c*_ along the virus length *x/L* obtained from simulations in Figs. 2C,F. The mean-field gradient length scale arising from cleaving is given by

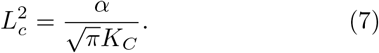

For the locomotion to occur, *L*_*c*_ must satisfy the condition 1 < *L*_*c*_ < *L*, where *L* is the non-dimensional virus length. Above the virus length *L*, the gradient is too shallow, rendering the binding landscape effectively uniform from the particle’s perspective. Below *l*_0_ the gradient is too steep, cleaving all receptor sites and detaching the virus from the substrate.

Figure 3 shows details of the viral locomotion as the function of the parameters: length *L*, NA catalytic activity *K*_*C*_, HA-SA dissociation constant *K*_*D*_, and friction Γ. The long-time MSD data for trajectories (e.g. Figs. 2B,E) is fitted to obtain the speed *v*. For both uniform and segregated HA/NA distributions, a minimal virus length *L* > *L*_*c*_ ≈ 10 is required to establish ballistic motion (Fig. 3A). While mean-field theory (dashed lines) predicts non-zero speeds for *L* < *L*_*c*_, the mean-field assumptions break down in this regime. As Γ*L* becomes larger, the speed *v* scales as *L*^−1^, in agreement with the analytical computations. Thus larger virions move more slowly through “sticky” environments.

**FIG. 3.**
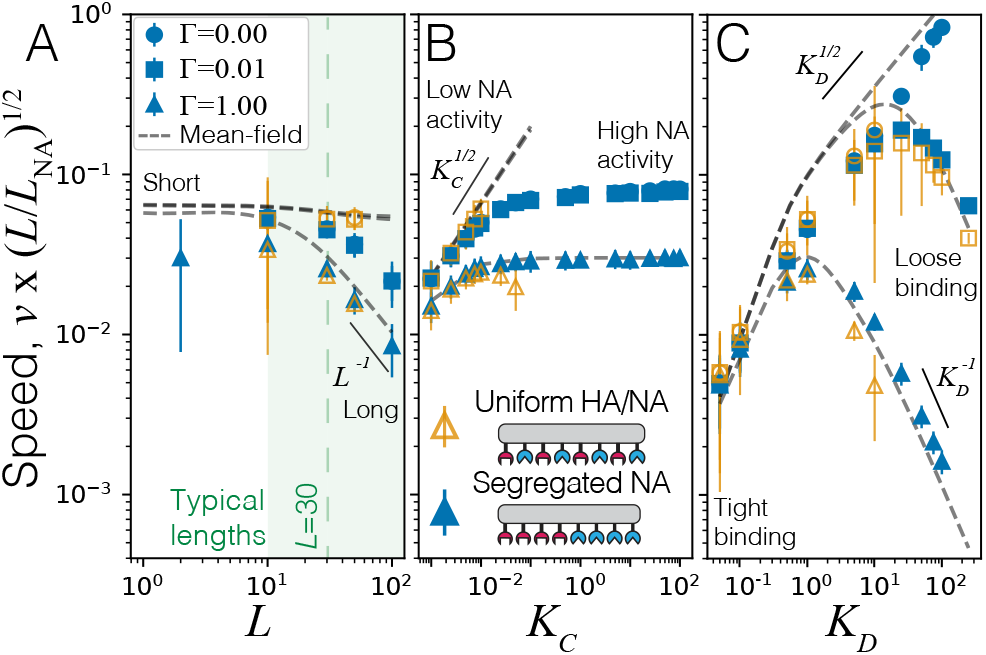
IAV motion varies with HA/NA activity and Γ. (A) *v* versus *L* shows SSB above a threshold *L* for the uniform case (orange), with *v* ~ *L*^−1^ for long *L*. Shaded region indicates typical IAV lengths; (*K*_C_= 0.075, *K*_D_ = 1). (B) *v* versus *K*_C_ indicates kinetic trapping beyond *K*_C_ ≈ 0.5 for a uniform distribution (orange), while the asymmetric case (black) exhibits a persistent *K*_C_-independent *v*; (*L* = 30, *K*_D_ = 1). (C) *v* versus *K*_D_ reveals an optimal *K*_D_, shifting right with decreasing Γ; (*L* = 30, *K*_C_ = 0.075). Markers denote different Γ values for uniform (orange) and asymmetric (black) cases; dashed lines are speeds from mean-field theory.

However, the role of the NA cleavage activity *K*_*C*_ (Fig. 3B) is strikingly different in the uniform and the segregated cases. For the uniform case, the predicted range for *K*_*C*_ from mean-field analysis is 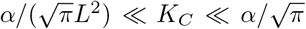, which for our parameters reads 10^−3^ ≪ *K*_*C*_ ≪ 0.5. Indeed, *v* abruptly drops to zero beyond a threshold *K*_*C*_ ≳ 0.5 (orange points), as excessive random cleaving prevents stable HA-SA binding, resulting in kinetic trapping (see SM for MSD plots). In contrast, the segregated case maintains robust motion even at high *K*_*C*_, approaching a constant terminal *v*. The physical restriction of NA prevents cleaving of all available SA sites, allowing HA to maintain binding without an upper limit on *K*_*C*_. The spatial segregation enforces a lower limit on *L*_*c*_, conferring resilience against fluctuations in NA activity. Discrepancies between theory and simulations arise from the mean-field approach’s heuristic addition of dissipation sources and its limitations in capturing local binding site depletion effects, e.g., when *K*_*C*_ ≳ 0.5.

Fig. 3C reveals a non-monotonic dependence of *v* on *K*_*D*_, with an optimal intermediate *K*_*D*_ maximizing speed. This defines a ‘Goldilocks’ constraint on *K*_*D*_, where both tight (low *K*_*D*_) and loose (high *K*_*D*_) binding hinder motion. At lower *K*_*D*_, increased effective friction results from longer bond lifetimes and higher average bond numbers [40, 41]. Conversely, at higher *K*_*D*_, HA-SA bonds generate insufficient driving force to maintain directed motion. We note that our model is more general compared to previous studies [26, 27], agreeing with them in the limit Γ*L* → 0 and reproducing the 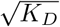 dependence at large *K*_*D*_. We solve the equation *∂v/∂K*_*D*_ = 0 to obtain the optimal value *K*_*D*,opt_. In the SM we calculate this derivative for two limiting cases: Γ*L* → 0 and Γ*L* → ∞. Interpolating between these cases we get the optimal *K*_*D*_ as:

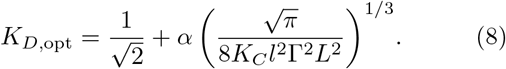

In the Γ*L* → 0 limit, the optimal *K*_*D*_ systematically increases as (Γ*L*)^−2*/*3^, going to ∞ as, in this regime, minimizing bond friction dominates. In the Γ*L* → ∞ limit, it goes to the constant value of 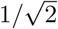. Thus, small changes in *K*_*D*_ or Γ*L* can give rise to markedly different transport speeds, whereas the catalytic rate *K*_*C*_ has a relatively minor effect.

Figure 4 maps the steady state speed in the parameter space of HA-SA dissociation constant (*K*_*D*_) and NA cleavage activity (*K*_*C*_). We choose Γ = 10^−2^ as physiologically relevant for IAVs navigating native mucinous environments, whose bulk viscosity can be much greater than that of water [42]. The heatmap shows different transport landscapes for uniform and polarized distributions. For the uniform case (Fig. 4A), effective movement is restricted to a narrow zone. In contrast, for the segregated case (Fig. 4B), an expansive high-speed regime spans a broad range of *K*_*D*_ and *K*_*C*_, indicating an inherent robustness conferred by asymmetric protein distribution on the viral envelope. We note that the 1 ~ nm/s speed is consistent with experimental findings [17].

**FIG. 4.**
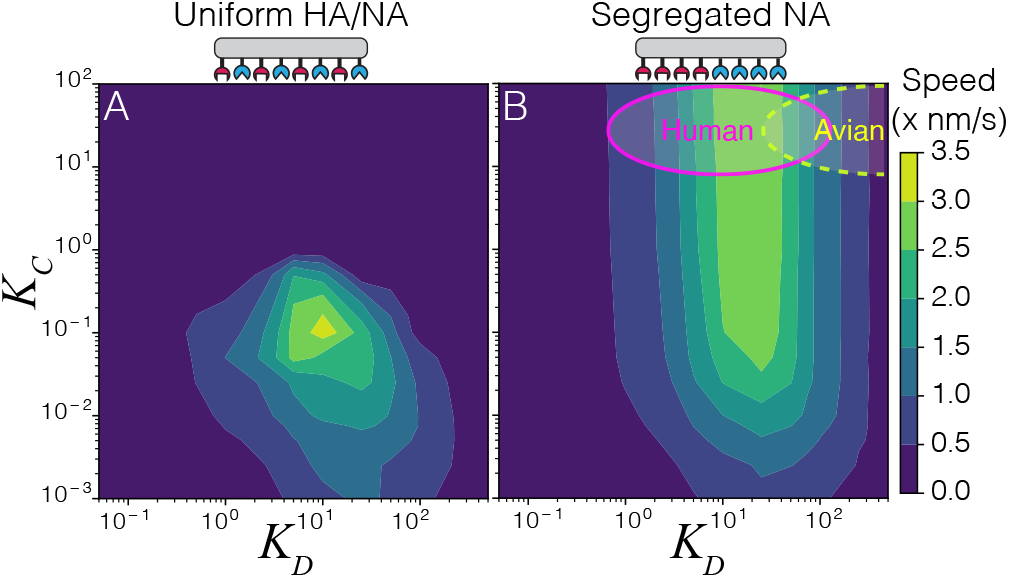
Speed heatmaps of IAV (*L* = 30, Γ = 0.01) as a function of HA-SA dissociation constant (*K*_D_) and NA-SA cleavage rate (*K*_C_). For uniform HA/NA distribution (A), motion is confined to a ‘sweet spot’. In contrast, the polarised case (B) exhibits an extended mobile regime, indicating spatial segregation broadens the ‘Goldilocks zone’ of optimal transport. Magenta outline: Range of *K*_D_ and *K*_C_ values for human IAV strains binding to/cleaving *α*2,6Sia in the human upper respiratory tract. Yellow dashed outline: Approximate *K*_D_ and *K*_C_ values for avian IAV strains binding to/cleaving *α*2,6Sia.

We show in Fig. 4B the estimated parameter ranges for human IAV strains binding to/catalyzing the *α*2,6-linked sialic acid (*α*2,6Sia) found in humans [43]. The measured *K*_*D*_ and *K*_*C*_ rates [31, 32, 44] lie within an optimal region (magenta) predicted by the model (see SM for details). While kinetic data for single HA-*α*2,6Sia bonds of avian IAV strains is lacking, the ~10x higher *K*_*D*_ value for *α*2,6Sia binding [45] corresponds to a distinct optimal cluster on the contour plot (yellow dashed outline), reflecting evolutionary adaptations to preferentially bind avian *α*2,3Sia receptors [46, 47]. Interestingly, retrospective analyses show historical pandemic precursor IAV strains acquired incremental HA mutations enhancing affinity for *α*2,6Sia receptors [48], a critical adaptation increasing human-to-human transmissibility. Furthermore, the presence of both *α*2,3Sia and *α*2,6Sia linkages in pigs allows them to serve as “mixing vessels” that facilitate IAV reassortment [49]. Thus, our framework provides a quantitative basis to understand how IAV receptor binding specificities contribute to host tropism and zoonotic transmission risk.

Our model also suggests that targeting IAV transport could be a useful antiviral strategy. While NA inhibitors (NAIs) like zanamivir and oseltamivir (Tamiflu) can reduce NA activity [50], our model shows that IAV locomotion is largely insensitive to NA activity. A dramatic shift along *K*_*C*_ axis is required to significantly block motion— which may contribute to the limited efficacy of NAIs in treating influenza [51, 52]. In contrast, we predict the HA-SA *K*_*D*_ to be a particularly attractive target for antivirals, as modest changes in on-rate or off-rate could more effectively move IAV out of the ‘Goldilocks zone’ required for directed motion.

## Conclusions

The rich phenomenological behavior of IAV locomotion stems from the interplay of stochastic receptor-ligand kinetics with dissipative mechanics governing motion in viscous environments. The existence of an optimal receptor-ligand binding-affinity range, finely tuned to the ambient viscosity, highlights a fundamental trade-off—overly tight binding hinders transport via excessive frictional forces, while loose binding forbids the transient receptor-ligand interactions required for motion. IAV locomotion hinges on a delicate balance between HA binding strength, NA cleavage activity, their spatial distribution on the virion surface, and the properties of the microenvironment. Notably, these equilibrium features of the virus-receptor system dictate even the out-of-equilibrium properties of viral motion. We find that segregating NA activity confers resilience, enabling robust directed movement across a broad range of receptor affinities and an even broader range of cleavage rates. The simulations and analytical theory yield remarkably consistent results, attributable to their shared fundamental assumptions. However, actual virus locomotion is more complex than our simplified model and further experimental data is crucial to validate our framework.

Our work identifies the molecular design principles important for a viral nanotransport strategy, insights that can be extended to many engineered systems [53–57]. Fundamentally, IAV-envelope organization appears engineered to leverage stochastic multivalent receptor-ligand kinetics for directed locomotion. The remarkable correspondence between the properties of IAVs and efficient transport across the epithelial barrier highlights this host defense as a critical evolutionary bottleneck. Viruses experience immense selective pressure to overcome multiple layers of host defenses and successfully infect target cells. Our findings show that locomotion through the mucociliary layer is an important source of evolutionary pressure. Since the optimal parameters differ across host species, mutations enabling zoonotic transmission must confer enhanced movement within the mucosal environment. Disrupting the balance between virion adhesive kinetics and mechanics may present opportunities for new antivirals that subvert this key aspect of influenza’s evolutionary prowess. Taken together, our findings could catalyze development of next-generation therapies specifically targeting influenza’s envelope organization and directed-locomotion strategy.

## Acknowledgements

We thank Guillaume Le Treut, David Yllanes, Amy Kistler, Michael Vahey, and members of the Fletcher lab for helpful discussions and suggestions. We also appreciate the constructive feedback from the anonymous reviewers. We thank CZ Biohub— SF and CZI Science for generous support. GH acknowledges a grant from CZI theory hub.

