## Supplemental Material for "Kinetics and optimality of influenza A virus locomotion"

### Supplemental Material to Kinetics and Optimality of influeza a virus locomotion

Siddhansh Agarwal and Daniel A. Fletcher
*Department of Bioengineering, University of California, Berkeley and*
*Chan Zuckerberg Biohub — San Francisco, 499 Illinois Street, San Francisco, CA 94158, USA*

Boris Veytsman
*Chan Zuckerberg Initiative, Redwood City, CA 94063, USA and*
*School of Systems Biology, George Mason University, Fairfax, VA 22030, USA*

Greg Huber
*Chan Zuckerberg Biohub — San Francisco, 499 Illinois Street, San Francisco, CA 94158, USA*
(Dated: August 9, 2024)

#### I. ANALYTICAL FORMALISM

Our calculations are based on the insight from the simulations that the most important processes are formation, breaking and cleavage of bonds between the virus and the substrate. The further insight is that the production of free energy due to irreversible bridge burning is dissipated through friction. Free energy of bonds is a log of the partition function. We assume a separation of time scales between the virus movement and the bond formation, so at any moment the configuration of bonds is close to the equilibrium one. We calculate equlibrium free energy $G$ as the function of the fraction of burned substrate sites $p_c$. Then we calculate the production of free energy due to increase of $p_c$, and equate it to the dissipation rate.

In the spirit of mean field approximation we neglect the details of interaction between the virus ligand sites, and assume that a substrate receptor site can participate in bonding with the probability $(1 - p_c)$. Then a virus site can be either free or bound to a substrate site at the distance $x$. The parition function for one virus site is

$$Z = 1 + \frac{1 - p_c}{K_D} \sum_x \exp(-x^2/\alpha^2) = 1 + \frac{(1 - p_c)n_{\mathrm{eff}}}{K_D}, \tag{S1}$$

where $n_{\mathrm{eff}} = \sum_x \exp(-x^2/\alpha^2)$ is is the effective number of substrate sites "seen" by a virus site, and the notation is given in Table I. This expression depends on the local value of $p_c$, which is different for different virus sites along the virus. The free energy is the sum of terms $-k_B T \log Z$ for each virus site. The summation can be changed to integration, giving

$$G = \frac{1}{l_0} \int_0^{Ll_0} \tilde{G}(x)\, dx, \quad \tilde{G} = -k_B T \log Z, \tag{S2}$$

where $L$ is the number of virus ligand sites.

The rate of burning is proportional to the probability that a substrate site is free. It also decreases as the distance from the virus site increases. We assume this decrease is described by the same factor $\exp(-x^2/\alpha^2)$ as the binding rate. After summation over all substrate sites, we get the burning rate at the given value of $p_c$ as

$$\frac{dp_c}{dt} = \frac{K_C(1 - p_c)n_{\mathrm{eff}} l}{l_0} \left[1 - \frac{\kappa}{K_D + (1 - p_c)n_{\mathrm{eff}}}\right] \tag{S3}$$

where

$$\kappa = \frac{1}{n_{\mathrm{eff}}} \sum_x \exp(-2x^2/\alpha^2). \tag{S4}$$

When there are many substrate sites per virus site, the summation can be changed to integration, which gives $n_{\mathrm{eff}} = \sqrt{\pi}\alpha l_0/l$ and $\kappa = 1/\sqrt{2}$.

The probability $p_c$ for a substrate site to be burned changes along the virus length $x$. In the stationary regime, when a virus moves with the constant speed $v$, it must be constant in the reference frame of the virus, which leads to the following PDE:

$$\frac{\partial p_c}{\partial t} - v \frac{\partial p_c}{\partial x} = 0, \tag{S5}$$

The solution to this equation with the boundary condition $p_c(0) = 0$ is

$$x = \frac{l_0 v}{K_C l n_{\text{eff}}(\kappa - K_D)} \left[ K_D \ln(1 - p_c) - \kappa \ln \left( 1 - \frac{p_c n_{\text{eff}}}{n_{\text{eff}} - \kappa + K_D} \right) \right]. \tag{S6}$$

This equation shows the different behavior of $p_c$ at the tail end of a very long virus ($x \to \infty$):

1. If $K_D < \kappa$, then at the tail end

$$\lim_{x \to \infty} p_c = 1 - \frac{\kappa - K_D}{n_{\text{eff}}}, \tag{S7}$$

and the characteristic length is

$$L_c = \frac{l_0 v \kappa}{K_C l n_{\text{eff}}(\kappa - K_D)}. \tag{S8}$$

2. If $K_D > \kappa$, then at the tail end

$$\lim_{x \to \infty} p_c = 1, \tag{S9}$$

and the characteristic length is

$$L_c = \frac{l_0 v K_D}{K_C l n_{\text{eff}}(K_D - \kappa)}. \tag{S10}$$

The total production of free energy $G$ is

$$\begin{aligned} \frac{dG}{dt} &= \frac{1}{l_0} \int_0^{Ll_0} \frac{d\tilde{G}(x)}{dt} dx = \frac{1}{l_0} \int_0^{Ll_0} \frac{d\tilde{G}(x)}{dp_c} \frac{\partial p_c}{\partial t} dx = \frac{v}{l_0} \int_0^{p_{c,\text{tail}}} \frac{dG(x)}{dp_c} dp_c \\ &= \frac{v}{l_0} \left[ \tilde{G}(p_{c,\text{tail}}) - \tilde{G}(p_c = 0) \right], \end{aligned} \tag{S11}$$

where $l_0$ is the distance between the ligand sites on the viral surface. Remarkably, the free energy production depends only on the difference in free energy between the head and the tail.

The production of free energy is dissipated through viscous damping and bond (or linker) friction. The first is proportional to the square of the speed $v$. We will write down the coefficient as $\gamma L l_0$, where $\gamma$ is the translational friction coefficient per virus site, reflecting the fact that for long filaments the friction is proportional to the length $L l_0$. For the linker friction calculations we follow the ideas of [1]. Namely, let us consider a substrate filament attached to a virus. The filament has the energy

$$U_{\text{el}} = \frac{k_B T x^2}{\alpha^2}, \tag{S12}$$

When the virus moves, this energy grows with the average rate

$$\left\langle \frac{dU_{\text{el}}}{dt} \right\rangle = \frac{2 k_B T v}{\alpha^2} \langle x \rangle, \tag{S13}$$

where averaging is performed over all filaments. If the lifetime of the bond is $\tau$, then before snapping the bond has the energy $k_B T v^2 \tau / \alpha^2$. This energy dissipates when the virus moves. Substituting the values of $\tau$ and $\alpha$ and summing over virus sites, we see that dissipation

$$\frac{dG}{dt} = \gamma L l_0 v^2 + \frac{S v^2}{2 k_{\text{off}} l_0} \int_0^{L l_0} p_b(x)\, dx. \tag{S14}$$

Choosing $l_0$ as our length scale and $k_{\text{on}}$ as inverse time scale, we get

$$\log \left[ \frac{K_D + n_{\text{eff}}}{K_D + (1 - p_c) n_{\text{eff}}} \right] = \frac{v}{\alpha^2} \left[ 2 \Gamma L + \frac{1}{K_D} \int_0^L p_b(x)\, dx \right] \tag{S15}$$

where, like in the main text, $\alpha^2 = 2k_BT/(Sl_0^2)$ and $\Gamma = \gamma k_{\rm on}/S$ Therefore,

$$v = \alpha^2 \frac{\log\left[(K_D + n_{\rm eff})\,/\,(K_D + (1-p_c)n_{\rm eff})\right]}{2\Gamma L + \int_0^L p_b(x)\,dx/K_D} \tag{S16}$$

To calculate the integral in this expression we note that $p_b$ is the function of $p_c(x)$:

$$p_b(x) = \frac{(1-p_c)n_{\rm eff}}{K_D + (1-p_c)n_{\rm eff}} \tag{S17}$$

Thus we change the integration over $x$ by the integration over $p_c$ using parametric equation (S6):

$$\begin{aligned} \frac{1}{K_D}\int_0^L p_b(x)\,dx &= \frac{1}{K_D}\int_0^{p_{c,\rm tail}} p_b(x)\frac{dx}{dp_c}dp_c \\ &= \frac{1}{K_D}\int_0^{p_{c,\rm tail}} \frac{v n_{\rm eff}\left(K_D + n_{\rm eff}(1-p_c)\right)}{v_0(1+K_D - n_{\rm eff}p_c)(K_D - \kappa + n_{\rm eff}(1-p_c))}dp_c \\ &= \frac{v}{v_0 K_D(1+\kappa - n_{\rm eff})}\left[(n_{\rm eff}-1)\log\left(1 - \frac{n_{\rm eff}\,p_{c,\rm tail}}{1+K_D}\right) - \kappa\log\left(1 - \frac{n_{\rm eff}\,p_{c,\rm tail}}{n_{\rm eff} - \kappa + K_D}\right)\right], \end{aligned} \tag{S18}$$

where $v_0 = K_C n_{\rm eff} l/l_0$. Fig. S1 plots the numerical solution as solid lines for $K_C = 0.075$, $L = 30$ for different $\Gamma$ values.

We now compute closed form expressions for the speed $v$ in some limits. We are interested in long viruses, i.e. $L \to \infty$. In this case equation (S6) shows that either i) $p_{c,\rm tail} = 1$, if $K_D > \kappa$; or ii) $p_{c,\rm tail} = 1 + (K_D - \kappa)/n_{\rm eff}$, if $K_D < \kappa$. In the first case, when $K_D > \kappa$,

$$\begin{aligned} \int_0^L p_b(x)dx/K_D &= \frac{v}{v_0 K_D(1+\kappa - n_{\rm eff})}\left[(n_{\rm eff}-1)\log\left(1 - \frac{n_{\rm eff}}{1+K_D}\right) - \kappa\log\left(1 - \frac{n_{\rm eff}}{n_{\rm eff}-\kappa+K_D}\right)\right] \\ &\approx \frac{v n_{\rm eff}}{v_0 K_D^2} + \ldots . \end{aligned} \tag{S19}$$

Similarly, in the second case, when $K_D < \kappa$, in the small $K_D$ limit, $p_b(x) \approx 1$

$$\int_0^L p_b(x)dx/K_D = L/K_D \tag{S20}$$

Equation (S16) then becomes

$$v = \begin{cases} \alpha^2 \dfrac{\log\left[(K_D + n_{\rm eff})\,/K_D\right]}{2\Gamma L + v n_{\rm eff}/(v_0 K_D^2)}, & K_D \gg \kappa \\ \alpha^2 \dfrac{\log\left[(K_D + n_{\rm eff})\,/\kappa\right]}{2\Gamma L + L/K_D}, & K_D \ll \kappa \end{cases} \tag{S21}$$

In the first case, when $K_D \gg \kappa$, this is a quadratic equation in $v$ with the solution

$$v = \frac{K_D^2 \Gamma L v_0}{n_{\rm eff}}\left(-1 + \sqrt{1 + \frac{n_{\rm eff}\alpha^2}{v_0\Gamma^2 L^2 K_D^2}\log\left[1 + n_{\rm eff}/K_D\right]}\right) \tag{S22}$$

In the case of negligible viscous friction, $\Gamma L = 0$, and

$$v = \alpha K_D \sqrt{v_0/n_{\rm eff}}\sqrt{\log(1 + n_{\rm eff}/K_D)} = \alpha\sqrt{v_0 K_D} + \ldots . \tag{S23}$$

In this case the speed *monotonically increases* with increasing $K_D$, independent of virus length $L$, and depends on the square root of the burning rate $K_C$ (through $v_0$).

Conversely, in the case of large friction, $\Gamma L \to \infty$,

$$v = \frac{\alpha^2}{2\Gamma L}\log(1 + n_{\rm eff}/K_D) = \frac{\alpha^2 n_{\rm eff}}{2\Gamma L K_D} + \ldots \tag{S24}$$

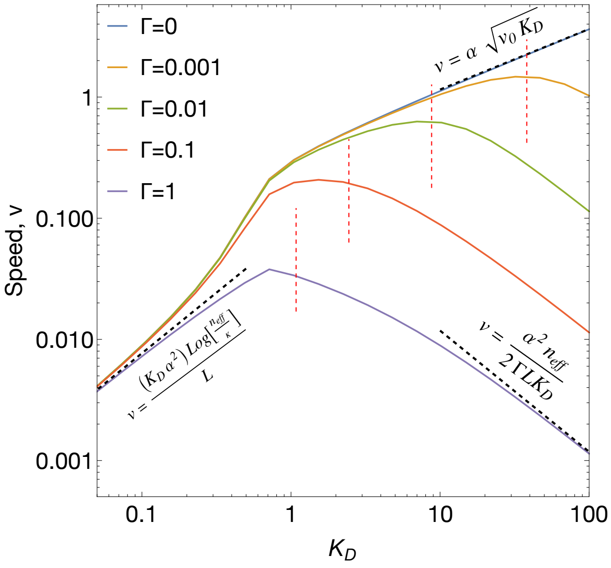


FIG. S1. Comparison of virus speed as a function of $K_D$ for various $\Gamma$ values, with fixed $K_C = 0.075$ and $L = 30$. Solid lines represent numerical solutions to the mean-field velocity, while red dashed lines indicate the optimal $K_D$ values predicted by eqn. (S28). Also plotted are the analytical expressions in various limits (black dashed lines) showing excellent agreement with the numerical solution.

The speed $v$ is independent of $K_C$ and *decreases monotonically* with increasing $K_D$, indicating a transition from the absence of an optimal $K_D$ when $\Gamma L = 0$ to the existence of an optimal value when $\Gamma L > 0$. Our model predicts a dramatic slowdown of motion ($v \sim 1/L$) for filament lengths $L \gg 10$ (Fig. 3A in main text). However, IAV particles exhibit a broad size distribution, with a small fraction of filaments reaching lengths up to $\sim 5\,\mu\text{m}$ [2]. This suggests that long filaments may serve other roles in the infection cycle, than the initial infection, potentially balancing reduced mobility with other functional advantages. The biological significance of this morphological diversity and its impact on IAV transmission dynamics warrants further investigation.

In the second case, when $K_D \ll \kappa$, this is a linear equation in $v$ with the solution

$$v = \frac{K_D \alpha^2}{L} \log\left[n_{\text{eff}}/\kappa\right]. \tag{S25}$$

The velocity increases linearly with $K_D$ and is independent of the friction $\Gamma$. Figure S1 plots as dashed black lines the closed form expressions (S23),(S24) and (S25), showing excellent agreement with the numerical solution.

##### A. Optimal $K_D$

Let us calculate the maximum of velocity (equation (S22)) as the function of $K_D$. At $K_D \gg n_{\text{eff}}$ we can write $\log(1 + n_{\text{eff}}/K_D) \approx n_{\text{eff}}/K_D$. Differentiating expression (S22) by $K_D$, we obtain for the maximal value of $K_D$ the expression

$$u + 4 - 4\sqrt{u+1} = 0, \quad u = \frac{n_{\text{eff}}^2 \alpha^2}{v_0 \Gamma^2 L^2 K_D^3}. \tag{S26}$$

This equation has the solution $u = 8$, which corresponds to

$$K_{D,\text{opt}} = \frac{n_{\text{eff}}^{2/3} \alpha^{2/3}}{2 v_0^{1/3} (\Gamma L)^{2/3}}. \tag{S27}$$

Similarly, in the limit $\Gamma L \to \infty$, we obtain $K_{D,\text{opt}} = \kappa$. Interpolating between the two limits we can write down

$$K_{D,\text{opt}} = \kappa + \frac{n_{\text{eff}}^{2/3} \alpha^{2/3}}{2 v_0^{1/3} (\Gamma L)^{2/3}}. \tag{S28}$$

Figure S1 presents a comparison between the results obtained from our simplified analytical expression (S28) (red dashed lines) and the numerical solutions to the mean-field velocity equation (solid lines). The graph demonstrates good agreement across a range of $\Gamma$ values, spanning from 0.001 to 1.

##### B. Range of validity of mean-field theory

In both $K_D \ll \kappa$ and $K_D \gg \kappa$ limits, the cleaving profile is exponential along the virus length, and $L_c = v/v_0$. The virus moves with constant speed as long as the condition $1 \ll L_c \ll L$ is satisfied. In the case of negligible viscous friction, $\Gamma L = 0$, and we obtain

$$L_c = \alpha\sqrt{\frac{1}{v_0}} = \sqrt{\frac{\alpha}{K_C\sqrt{\pi}}}. \tag{S29}$$

The limits on the cleavage rate are therefore,

$$\frac{\alpha}{\sqrt{\pi}L^2} \ll K_C \ll \frac{\alpha}{\sqrt{\pi}} \tag{S30}$$

Conversely, in the large $\Gamma L$ case,

$$L_c = \frac{\alpha^2}{\Gamma L v_0} = \frac{\alpha}{L\Gamma K_C\sqrt{\pi}}, \tag{S31}$$

and the bounds on the cleavage rate read,

$$\frac{\alpha}{\sqrt{\pi}L^2\Gamma} \ll K_C \ll \frac{\alpha}{\sqrt{\pi}L\Gamma}. \tag{S32}$$

##### C. Power balance vs. force balance approach

While a force balance approach is often employed (cf. [3]), our analysis utilizes a dissipative power balance method. This eliminates the need for explicit calculations of individual forces acting on the virus (e.g., viscous drag, bond tensions) and offers a simplicity and clarity while yielding equivalent insights. To illustrate the connection between these methods, we can derive the force acting on a virus from power dissipation consideration. Divding the power production by speed $v$ yields the propulsive force $F$:

$$F \sim \frac{1}{v}\frac{dG}{dt} = \frac{1}{l_0}\left[\tilde{G}(p_{c,\mathrm{tail}}) - \tilde{G}(p_c = 0)\right]. \tag{S33}$$

Notably, the force depends solely on the free energy production difference between the virus head and tail and results in:

$$F = \frac{k_B T}{l_0}\log\left[\frac{K_D + n_{\mathrm{eff}}}{K_D + (1-p_c)n_{\mathrm{eff}}}\right]. \tag{S34}$$

In the regime where $K_D \gg 1/\sqrt{2}$, we can approximate $p_{c,\mathrm{tail}} \approx 1$, simplifying the equation to:

$$F \sim \frac{k_B T}{l_0}\log\left[1 + \frac{n_{\mathrm{eff}}}{K_D}\right]. \tag{S35}$$

For the typical values of parameters $F \sim 0.1$–$1\,\mathrm{pN}$.

#### II. STOCHASTIC SIMULATION IMPLEMENTATION

We use an overdamped equation of motion for the dynamics of the center of mass of the simulated virus $x(t)$, excluding explicit thermal noise:

$$v = -\frac{1}{\Gamma L}\sum_i \Delta x_{ij}, \quad v = dx(t)/dt, \tag{S36}$$

where $\Delta x_{ij}$ is the distance between the $i$th ligand on the viral surface bound and the $j$th receptor on the substrate. $\Gamma = \gamma k_{\rm on}/S$ is the ratio of spring relaxation to bond formation timescales and represents friction coefficient per unit length. At each time step, we update the configuration of bound springs and cleaved receptors via the Gillespie algorithm [4, 5]. Briefly, the dimensionless binding/unbinding rates are given by transition-state theory as:

$$q_{\rm off} = \frac{K_D}{1+\exp(-\Delta x_{ij}^2/\alpha^2)}, \quad q_{\rm on} = \frac{\exp(-\Delta x_{ij}^2/\alpha^2)}{1+\exp(-\Delta x_{ij}^2/\alpha^2)}. \tag{S37}$$

This defines an effective stretch-dependent dissociation constant $q_{\rm off}/q_{\rm on} = K_D \exp(\Delta x/\alpha^2)$, where $K_D = k_{\rm off}/k_{\rm on}$. The dimensionless NA cleavage rate is:

$$q_c = K_C \exp(-\Delta x/\alpha^2), \tag{S38}$$

where $K_C = k_c/k_{\rm on}$. Each ligand/receptor is treated as a distinct chemical species as the binding/unbinding/cleaving rates are configuration dependent. In particular, the algorithm evolves the number of bonds and also tracks the list of ligand/receptor molecules that form a bond.

We sample the list of reactions and the time for them to happen ($\tau_\ell, \ell = 1, 2, \ldots$). To sample a reaction event, let $\mathcal{A} = \{a_{11}^X, a_{12}^X, \ldots, a_{1N_R}^X, \ldots, a_{N_L 1}^X, a_{N_L 2}^X, \ldots, a_{N_L N_R}^X\}$ be the array containing all possible ligand-receptor pairs, where $a_{ij}^X$ is the propensity for ligand $i$ and receptor $j$ to have a reaction of type $X$ ($X$ = on/off for binding/unbinding and $X$ = cleave for enzymatic cleavage). Thus, $a_{ij}^{\rm on} = q_{\rm on}$ if ligand $i$ and receptor $j$ can bind, $a_{ij}^{\rm off} = q_{\rm off}$ if already bound and $a_{ij}^{\rm cleave} = q_c$ if ligand $i$ can irreversibly cleave receptor $j$, 0 otherwise. The total propensity is $a_T = \sum_{i=1}^{N_L} \sum_{j=1}^{N_R} a_{ij}^X$. Given $a_T$, we sample $\tau_\ell$ from a Poisson distribution with the average $(a_T)^{-1}$.

Then the net instantaneous force is computed and used to update the speed and position of the virus by using the analytical solution to (S36),

$$x_{\rm new} = \frac{1}{N_B}\sum_i \Delta x_{ij} + \exp(-N_B \tau_\ell/(\Gamma L))\left(x_{\rm old} - \frac{1}{N_B}\sum_i \Delta x_{ij}\right), \tag{S39}$$

where $N_B$ is the number of bound ligand sites on the virus at that time instant. Note that in the limit $\Gamma \to 0$, the system dynamics are governed only by the first term on the RHS.

We then select an event with probability $\mathbb{P}(j) = a_{ij}^X/a_T$, choosing the $i$th ligand and $j$th receptor pair and implementing the corresponding reaction of type $X$. Before sampling the next step, we update the affinity matrix (along with $a_T$).

For each set of parameters, we ran N = 64 independent trajectories with randomized initial conditions up to a terminal time $T = 10^4$. Trajectories were propagated using an analytical solution to Eq. S36 coupled to the Gillespie algorithm to update bound ligand-receptor pairs and cleaved receptors. From each trajectory, we computed the time-averaged mean-squared displacement (MSD):

$$\langle \Delta r^2(t)\rangle_T = \frac{1}{T-t}\int_0^{T-t} [r(t'+t) - r(t')]^2\, dt', \tag{S40}$$

where $r(t)$ is the particle position at lag-time $t$ and $T$ is the total trajectory duration. The MSD quantifies the average particle displacement over a lag time $t$. We averaged the individual MSD curves over the N = 64 trajectory ensemble to obtain the final ensemble-averaged MSD $\langle \Delta r^2(t)\rangle$, plotted in Figure 2 of the main text. In the steady-state, the ensemble-averaged MSD and the MSD obtained by time averaging the individual $\Delta r^2(t)$ curves first are equivalent. For times $t$ long compared to the relaxation time, the ensemble MSD grows as $\langle \Delta r^2(t)\rangle \sim v^2 t^2$, where $v$ is the long-time ballistic speed. We extracted $v$ by fitting $\ln\langle \Delta r^2(t)\rangle$ vs $\ln t$ to a line with slope 2 over the last 2000 time units. Error bars in $v$ have contributions from the fitting procedure and statistical uncertainty from the finite ensemble size N. The fitting uncertainty was estimated from the covariance matrix. The statistical uncertainty was the standard deviation of the N individually fitted speeds about their mean.

The MSD curves exhibit a sequence of dynamical regimes. At very short times the MSD scales ballistically, reflecting virus motion before bonds form. As intermediate times, bonds form and relax, and the dynamics transition to a diffusive regime characteristic of systems with dynamical constraints or intermittent trapping from the transient binding/unbinding kinetics. Remarkably, at long times $t \gg \tau_c$, where $\tau_c$ is the timescale for substantial receptor cleavage, the MSD curves develop an emergent superdiffusive ballistic regime $\langle \Delta r^2(t)\rangle \sim t^\alpha$, $\alpha = 2$. This regime arises from the irreversible cleaved of receptors generating free energy, causing the virus to behave as an active particle with self-propelled motion away from its cleaved trail.

#### III. PARAMETERS ESTIMATION

Let us estimate model parameter values for a typical influenza A virion. IAVs have an HA density of 20 000 µm$^{-2}$ $\sim$ 0.02 nm$^{-2}$. Assuming these are distributed uniformly along the rectangular cross-section presented to the substrate, the average spacing between points is $l_0 \sim 1/\sqrt{0.02} \sim 10\,\mathrm{nm}$. The substrate (SA) density is typically of the same order (or denser) than the HA density; hence $l/l_0 \approx 0.1 - 1$ (we choose $l/l_0 = 0.25$). The number of substrate sites a given virus site "sees" $n_{\mathrm{eff}}$ depends on the ratio $\alpha/l$, where $\alpha = \sqrt{2k_BT/(Sl_0^2)}$. Typically, the spring constant $S \sim 0.01$–$1\,\mathrm{k_B T/nm^2}$ (we choose $S = 0.02\,\mathrm{k_BT/nm^2}$ so that $\alpha = 1$), therefore $n_{\mathrm{eff}} = \sqrt{\pi}\alpha/l \approx 10$. $\kappa = (1/n_{\mathrm{eff}})\sum_x \exp\left(-2x^2/\alpha^2\right) = 1/\sqrt{2}$ for $\alpha/l \gtrsim 1$.

In order to estimate the friction coefficient per unit ligand $\gamma$, we use the Stokes drag on a sphere so that $\gamma = 6\pi\eta l_0$, where $\eta$ is the dynamic viscosity of the surrounding mucus environment $10^{-1}$–$10^2\,\mathrm{Pa\cdot s}$ [6], many orders of magnitude larger than that of water. Inserting this into the Stokes drag formula, we obtain $\gamma \approx 10^{-6}$–$10^{-9}$ kg/s as a rough estimate. Alternatively, using Brenner theory [7] for dilute suspensions of cylindrical rods with lengths $L(\approx 300\,\mathrm{nm})$ and diameters $d(\approx 50\,\mathrm{nm})$, we obtain $\gamma = (2\pi\eta L/\log(2L/d - 0.5))/30 \approx 10^{-6}$–$10^{-9}$ kg/s, consistent with the simple Stokes drag estimate. Using these estimates, we can compute the timescale for a passive $\sim 300\,\mathrm{nm}$ particle to diffuse through the viscous $\sim 1 - 10\mu\mathrm{m}$ thick mucus layer to be $t_d \sim \sqrt{10\times10^{-6}/(k_BT/(30\times10^{-9}))} \sim 2\,\mathrm{h}$, in agreement with [8].

We estimate $K_D$ using the estimates from [8]. In our theory, we define the native $K_D = k_{\mathrm{off}}/k_{\mathrm{on}}$ at zero bond extension. Here, $k_{\mathrm{on}}$ and $k_{\mathrm{off}}$ are probabilities per unit time. In contrast, experimental measurements, are in terms of the bulk on-rate $k_{\mathrm{on,b}} \approx 150$–$1000\,\mathrm{M^{-1}s^{-1}}$ [9, 10] and the off-rate $k_{\mathrm{off}} \approx 30\,\mathrm{s^{-1}}$, for a single HA-SA bond [11]. In order to go from these units to probabilities we need a characteristic interaction length scale $b$. We use $b \sim 7.5\,\mathrm{nm}$ [8, 12], modeling the flexibility of the carbohydrate and protein backbone to which sialic acid is attached, along with the potential flexibility of the HA/NA on the virus surface. Thus, we obtain

$$k_{\mathrm{on}} = \frac{3k_{\mathrm{on,b}}}{(4/3)\pi 1000 b^3 N_{\mathrm{Av}}} \approx 0.4\text{–}2\,\mathrm{s^{-1}}. \tag{S41}$$

This results in a $K_D \approx 10 - 100$. The extra factor of 3 in the denominator accounts for the trimeric nature of HA: each HA can bind to 3 SA moieties. Since the concentration of sialic acid is high relative to the $K_M$ for NA ($> 10\,\mathrm{mM}$ versus 100–1000 mM [9]), the burning rate of SA by NA is determined from the catalytic rate constant (and does not depend on $K_M$). Hence, $k_c \approx 10$–$100\,\mathrm{s^{-1}}$, or $K_C \approx 10$–$100$ (choosing $k_{\mathrm{on}} = 1\,\mathrm{s^{-1}}$).

The friction coefficient, $\Gamma = \gamma k_{\mathrm{on}}/S$ can span a wide range from $10^{-5}$–$10^2$, depending on the environment; in the main text we choose $\Gamma \sim 0.01$ corresponding to $\eta \sim 50\,\mathrm{Pa\cdot s}$, $k_{\mathrm{on}} = 1\,\mathrm{s^{-1}}$ and $S = 0.02\,\mathrm{k_BT/nm^2}$.

#### IV. TABLE OF PARAMETERS AND THEIR VALUES

TABLE I: Parameters and their values

| Parameter | Description | Range of values | Reference(s) |
|---|---|---|---|
| **Dimensional parameters** | | | |
| $L'$ | IAV length | 100 nm–5 µm | [13, 14] |
| $L_m$ | Mucous layer thickness | 1–10 µm | [15–17] |
| $S$ | HA-SA bond spring constant | 0.01–1 $\mathrm{k_BT/nm^2}$ | [3] |
| $\eta$ | Bulk viscosity of mucus | $10^{-1}$–$10^2$ Pa · s | [6] |
| $\gamma$ | Translational friction coefficient per ligand | $10^{-6}$–$10^{-9}$ kg/s | |
| $k_c$ | NA-SA catalytic rate | 10–100 $\mathrm{s^{-1}}$ | [9] |
| $k_{\mathrm{off}}$ | HA-SA off rate | 30 $\mathrm{s^{-1}}$ | [11] |
| $k_{\mathrm{on}}$ | HA-SA on rate | 0.4–2 $\mathrm{s^{-1}}$ | [9, 10] |
| $l'$ | SA receptor spacing | 2.5 nm | [8] |
| $l_0$ | Ligand (HA) size | 10 nm | [18] |
| $t_c$ | Mucociliary clearance timescale | 20 min | [8] |
| $t_d = \sqrt{L_m/(k_BT/\gamma L)}$ | Passive diffusion timescale | 1–74 h | [8] |
| **Dimensionless parameters** | | | |
| $\alpha = \sqrt{2k_BT/(Sl_0^2)}$ | Thermal fluctuation parameter | 1 | |
| $\Gamma = \gamma k_{\mathrm{on}}/S$ | Friction parameter | 0, 0.01 and 1 | |

TABLE I: Parameters and their values, continued

| Parameter | Description | Range of values | Reference(s) |
|---|---|---|---|
| $K_C = k_c/k_{on}$ | NA-SA catalytic rate | 0.0001–100 | |
| $K_D = k_{off}/k_{on}$ | HA-SA dissociation constant | 0.005–100 | |
| $L = L'/l_0$ | IAV length | 30 | |
| $l = l'/l_0$ | SA receptor spacing | 0.25 | |

#### V. COMPARING UNIFORM VS. SEGREGATED NA CONFIGURATIONS AS $K_C$ INCREASES

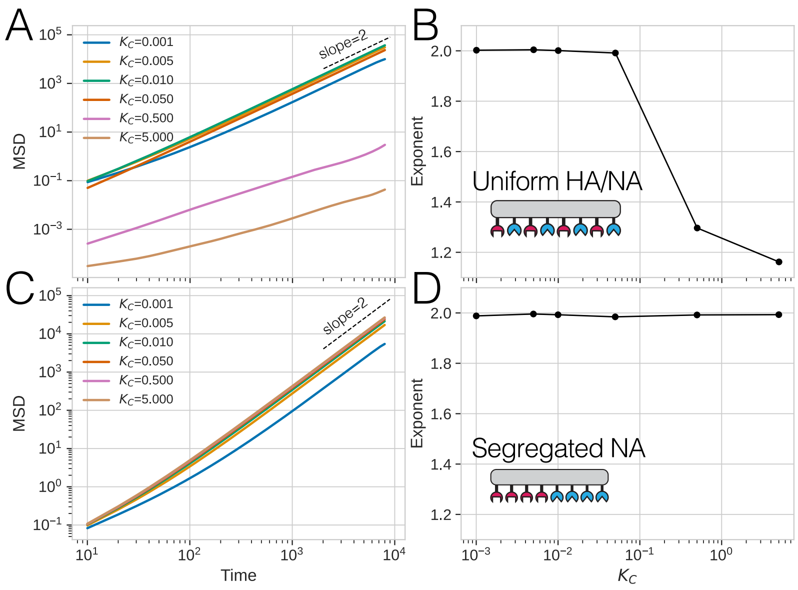


FIG. S2. (A, C) Mean square displacement (MSD) vs time and (B, D) long-time exponent as a function of cleaving rate $K_C$ for uniform (top panel) and segregated NA configurations (bottom panel) of IAV for $K_D = 1$ and $L = 30$.

Figure S2 illustrates the mean square displacement (MSD) behavior derived from simulations of IAV particles with varying HA and NA surface configurations as a function of increasing cleaving rate $K_C$. These plots help to explain the phenomena described in Fig. 3B of the main text. We present a comparative analysis of uniform distributions versus segregated NA configurations, with fixed parameters of $K_D = 1$ and $L = 30$. As $K_C$ increases, we observe a distinct transition to a kinetically trapped state at approximately $K_C \approx 0.5$ in the uniform configuration case. This transition is manifested by a significant decrease in the long-time MSD slope, as depicted in Fig. S2B. In contrast, the segregated NA configuration (Fig. S2D) exhibits remarkable robustness to variations in $K_C$, maintaining a ballistic exponent (slope = 2) across the entire range of cleaving rates examined.

#### VI. NOTE ON IAV TRANSPORT THROUGH MUCUS

Virus transport through mucus is a complex process with limited available experimental data. We consider a mucus layer thickness range of 1–10 µm and assume a typical virus velocity of $v_0 = 3$ nm/s (from Fig. 4B in the main text). For purely ballistic motion, the transit time across the mucus layer would range from $\sim 5.6$ min (for the thickness 1 µm) to $\sim 55.6$ min (for the thickness 10 µm). However, the actual transport mechanism likely combines ballistic motion and diffusion. The effective diffusion coefficient can be expressed as [19]: $D_{\text{eff}} = D_0 + v_0^2\tau/d$ where $D_0 \approx 10^{-4}$ µm$^2$/s is the passive diffusion coefficient, $\tau$ is the angular decorrelation time, and $d = 3$ for three dimensional motion. The second term depends on $\tau$. For $\tau \approx 1$ ms it is $3 \times 10^{-9}$ µm$^2$/s, while for $\tau \sim 1000$ s it is $3 \times 10^{-3}$ µm$^2$/s, and dominates the first term.

A Brownian particle crossing a 1–10 µm layer in 20 minutes needs to have a diffusion coefficient $D_0 \approx 10^{-4}$–$10^{-2}$ µm$^2$/s. This shows the necessity of some degree of sustained directional motion. The exact transport regime (ballistic, diffusive, or combined) critically depends on $\tau$, which is challenging to estimate theoretically due to complex mucus rheology and virus-mucus interactions. These complexities arise from mucus's non-Newtonian properties, heterogeneous composition, and potential specific biochemical interactions between viral surface proteins and mucus components [6, 20]. Additional barriers to virus transport, such as the glycocalyx and other cellular structures, further complicate these estimates. Further research combining theoretical modeling with advanced experimental techniques is needed to probe the dynamics at relevant spatial and temporal scales and address these questions more comprehensively.

---
